# Komenti: A semantic text mining framework

**DOI:** 10.1101/2020.08.04.233049

**Authors:** Luke T Slater, William Bradlow, Robert Hoehndorf, Dino FA Motti, Simon Ball, Georgios V Gkoutos

**Affiliations:** Institute of Cancer and Genomic Sciences, University of Birmingham, UK; University Hospitals Birmingham, NHS Foundation Trust, United Kingdom; Computational Bioscience Research Center, King Abdullah University of Science and Technology, SA; MRC Health Data Research UK (HDR UK) Midlands, UK

## Abstract

**Summary:** Komenti is a reasoner-enabled semantic query and information extraction framework. It is the only text mining tool that enables querying inferred knowledge from biomedical ontologies. It also contains multiple novel components for vocabulary construction and context disambiguation, which can improve the power of text mining and ontology-based analysis tasks, with a view towards making full use of the semantic provision of biomedical ontologies for text characterisation and analysis. Here, we describe Komenti and its features, and present a use case wherein we automate a clinical audit, extracting medications for hypertrophic cardiomyopathy patients from text, revealing a high precision, and identifying a sub-cohort of patients with atrial fibrillation who are not anti-coagulated, and are therefore at a higher risk of stroke.

**Availability and Implementation:** Komenti is freely available under an open source licence at http://github.com/reality/komenti.

More information concerning the use-case is available in supplementary data.

## Introduction

Biomedical ontologies contain a wealth of information in the form of natural language labels and synonyms associated with terms, together forming controlled domain vocabularies that describe entities within scientific domains. As a direct consequence, ontologies form a powerful resource for text mining [1]. While text mining has been an important tool in biomedicine for a long time, it is becoming increasingly vital in a clinical setting, where text extraction can be used to aid audits, service improvement, and research, without a reliance on time-consuming and burdensome manual data extraction, which has historically been performed by clinical teams themselves. Frameworks such as CogStack [2] make use of medical domain vocabularies, and have recently shown promising applications to clinical text mining, revealing the ability to characterise patients and diseases with a power previously unreported.

However, the features of biomedical ontologies exceed the provision of controlled domain vocabularies [3]. They also provide a rich semantic underpinning by means of description logics, enabling semantic information retrieval and analysis. Recent works in ontology-based machine learning and vectorisation, such as OPA2Vec [4], are bridging the gap between text mining, machine learning, and semantic analysis, and they highlight the need for novel approaches that make full use of the descriptive and semantic features provided by biomedical ontologies [5].

We developed Komenti, a semantic text mining framework that exploits the expressive semantic features of biomedical ontologies, facilitating access to knowledge from their totality. Moreover, Komenti includes several novel text mining components, which have been shown to outperform existing methods and improve analyses. By leveraging novel technologies in both of these areas, with tight integration, Komenti presents a promising new framework for semantic text characterisation and analysis.

## Approach

The Komenti framework provides a set of tools for information retrieval and extraction, including advanced semantic query mechanisms, biomedical corpus retrieval, named entity recognition, context disambiguation, and analysis, including stratification of clinical observations. These tools can either be used in isolation or together as part of an information extraction pipeline. The framework, which is written in Groovy, employs background biomedical knowledge combined with text mining approaches, and can be applied directly to clinical, biomedical, or literature text to facilitate insights into the concepts and relationships they describe. Below, we describe the main functionalities of the framework, how they interact, and can be applied for analysis. Finally, we provide an example of its application in performing a clinical audit of hypertrophic cardiomyopathy patient medications.

The query commands interface with the AberOWL reasoner-based ontology access system [6], enabling direct access to inferred knowledge from ontologies. There are no existing command line tools to easily obtain classes and labels via the semantic features of biomedical ontologies. For example, in a single command Komenti can build a text mining vocabulary from the parent term `Medical Intervention (0AE:0000002)` in the COVID-19 ontology [7], including labels surrounding drug development, fluid management, and medical examinations relevant to the COVID crisis. This notion can also be extended to complex class expressions, returning terms that match a given axiomatic description, such as `‘part of’ some apoptosis`.

Komenti also includes a novel vocabulary expansion algorithm, adding additional labels and synonyms for terms that can be matched in text, by linking equivalent classes between ontologies using lexical and semantic queries. This has provisionally been shown to vastly increase the scale of vocabulary available in several ontologies, the amount of information returned in information retrieval tasks, and to improve the performance of semantic analysis of clinical text [8].

Komenti can directly obtain Pubmed literature abstracts, with a number of options for disjunctive and conjunctive query based on groups of vocabulary terms. Thereby, a relevant corpus with which to perform text analysis can be directly obtained. Since these results can be saved as plain text, they also enable external analysis of texts, or use of Komenti as an aid for semi-automated or manual literature review.

For information extraction, Komenti implements Stanford CoreNLP [9], pre-processing text with its tokenization and sentence splitting annotators. By passing the vocabulary’s labels and synonyms into its RegexNER annotator, it provides a fast lexical named entity recognition system for ontology terms. An Apache PDFBox implementation caters the direct annotation of PDF files. For context disambiguation, Komenti uses a novel negation detection algorithm, which has shown improved stability and performance over several popular solutions in tests over clinical data in two settings [10]. It also contains a variation of the negation detection algorithm, using configurable uncertainty vocabulary to determine whether concepts are uncertain in a particular context (e.g. a patient is being tested for a disease). Sentences can also be excluded from annotation via a configurable list of string phrases, or labelled as concerning a family member. Latterly, information concerning family history of clinical findings can be discerned, as well as exclusion of references to the patient.

Since Komenti outputs annotations in a simple tabular format, analysis software can easily make use of the produced information. In a previous experiment, labels derived by Komenti from clinical text were used in semantic similarity analyses [8]. Komenti also provides several features for internal analyses of its annotations. For example, the *diagnose* command uses a count-based decision procedure, based on annotation contexts (e.g. negation, uncertainty), to classify a bearer’s status for findings described by annotations (e.g., does a patient suffer from a disease, or are they under a medication?). Komenti can also analyse annotations to report co-occurrence of entities, and to suggest logical axioms. While many of these features are in development, they demonstrate that Komenti is a robust basis for the development of experimental and analytic tools. Komenti also provides configurable templates for sequential execution of its tools. This feature turns it from a collection of related commands into an integrated framework for designing, executing, and sharing semantically enabled text analysis pipelines.

## Use Case: Medication Audit

We used Komenti to extract and classify anticoagulant medications from text records for a population of patients with hypertrophic cardiomyopathy (HCM) at University Hospitals Birmingham NHS Trust. Using the patient characterisation workflow described in the Komenti documentation, it determined which patients were on anticoagulant medications. A manual validation of 52 patients by a clinical specialist, including affirmed, uncertain, and negative cases, revealed a precision of **0.959.** The approach also identified 39 patients with HCM and atrial fibrillation, who were not being treated with anti-coagulant medication, leaving them at increased risk of stroke, and who will now be followed up by the specialist service. More information about the use case methods and results are given in supplementary materials.

## Conclusion

Komenti tightly integrates semantically enabled ontology access and text mining, introducing several novel components. It integrates background biomedical knowledge and facilitates its use on any biomedical text, providing a framework for information retrieval, extraction, and analysis. NLP is playing an increasingly important role in clinical settings, wherein automated approaches promise to accelerate the evaluation of care quality across large patient groups, drive rapid service improvement, and improve patient/disease stratification. Our use case highlighted Komenti’s effectiveness for this purpose by identifying patients receiving suboptimal treatment with accuracy equivalent to that of an expert.

## Ethics and Funding

The UoB Ethical Review approved this work (ERN_20-0338). GVG and LTS acknowledge support from support from the NIHR Birmingham ECMC, the NIHR Birmingham SRMRC, Nanocommons H2020-EU (731032), and the NIHR Birmingham Biomedical Research Centre and the MRC HDR UK (HDRUK/CFC/01).

## Supplementary Material

### Use Case: Materials and Methods

A guide describing how Komenti can be used to characterise patients and associated concepts is online at https://github.com/reality/komentLguide/blob/master/patients.md. This experiment was performed using an earlier prototype of the Komenti software with a different user interface, although all core functionalities were equivalent.

A vocabulary was constructed through a collaboration between a clinical expert (WB) and the researcher (LTS), wherein candidate drugs were identified, and linked to ontology terms. Komenti’s query feature was then used to obtain a labels and synonyms for the drugs to be matched in text. Patients with HCM and atrial fibrillation were previously identified in another audit. Clinical letters discussing those patients were obtained from the document server at the University Hospitals Birmingham NHS Trust on 02/06/2019. The annotation and diagnose tools were then used on the documents to identify which patients were prescribed anticoagulants. Each patient received a classification of either affirmed, negated, uncertain, or unmentioned for each drug. Unmentioned classifications were discarded from further analysis. Validation was performed manually on mentioned classifications for 52 patients sampled from the results by the clinical specialist (WB), based on the sentence contexts provided by the tool, and evaluation of the overall context of the document the mentions appeared in. Recall was not calculated, because there was no available gold standard, and the validation only recorded binary results for assertions, not which label it should have been given.

The University of Birmingham Ethical Review approved this work (ERN_20-0338). No informed consent was required as this was a service improvement project, and the documents were not de-identified as we intend to follow up individuals lost to discharge. The hospital identification number linked documents belonging to the same patient and associated data to the registry[8]. Data remained within the Trust and only included information related to HCM.

### Use Case: Results

**Table 1.** Number of patients in each group. The object of the study was to identify patients who had atrial fibrillation, but were not taking an anti-coagulant.

| **Group** | **Count** |
| --- | --- |
| Patients | 822 |
| Patients with AF | 115 |
| Patients with AF and no anti-coagulant | 39 |

**Table 2.**
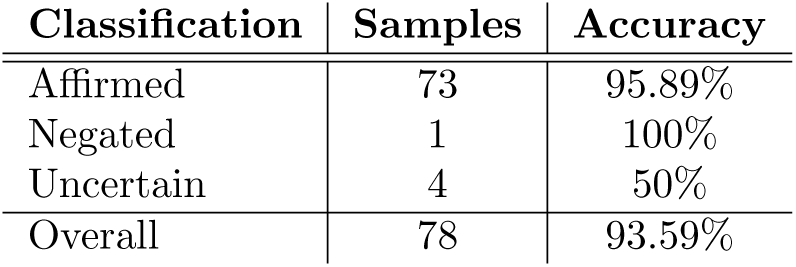
Overall accuracy for each overall classification category. Due to the small sample sizes, this doesn’t form a robust evaluation of the performance of the negation and uncertainty algorithms, but helps hilight how they are used to exclude patients from the affirmed group. The overall precision was **0.959**.

| **Classification** | **Samples** | **Accuracy** |
| --- | --- | --- |
| Affirmed | 73 | 95.89% |
| Negated | 1 | 100% |
| Uncertain | 4 | 50% |
| Overall | 78 | 93.59% |

**Table 3.** Per-class accuracy of assertions over the 52 patients manually validated. Assertions with no samples are omitted.

| **Drug** | **Status** | **Accuracy** | **Samples** |
| --- | --- | --- | --- |
| ramipril | Affirmed | 95.24% | 21 |
|  | Uncertain | 100% | 1 |
| apixaban | Affirmed | 100% | 4 |
| warfarin | Negated (family history) | 100% | 1 |
|  | Affirmed | 100% | 17 |
|  | Uncertain | 0% | 1 |
| spironolactone | Affirmed | 85.71% | 7 |
|  | Uncertain | 100% | 1 |
| lisinopril | Affirmed | 100% | 1 |
| losartan | Affirmed | 100% | 7 |
|  | Uncertain | 0% | 1 |
| candesartan | Affirmed | 87.5% | 8 |
| rivaroxaban | Affirmed | 100% | 4 |
| dabigatran | Affirmed | 100% | 1 |
| eplerone | Affirmed | 100% | 1 |
| ivabradine | Affirmed | 100% | 1 |
| edoxaban | Affirmed | 100% | 1 |

